# Glutamatergic and cholinergic metabotropic modulation induces plateau potentials in hippocampal OLM interneurons

**DOI:** 10.1101/508382

**Authors:** Nicholas Hagger-Vaughan, Johan F. Storm

**Affiliations:** Section for Physiology, Institute of Basic Medical Sciences, University of Oslo, Oslo, Norway

## Abstract

Oriens-lacunosum moleculare (OLM) cells are hippocampal inhibitory interneurons that have been implicated in regulation of information flow and synaptic plasticity in the CA1 circuit.

Since anatomical evidence indicate that OLM cells express metabotopic cholinergic (mAChR) and glutamatergic (mGluR) receptors, such modulation of these cells may contribute to switching between functional modes of the hippocampus.

Using a transgenic mouse line to identify the Chrna2-positive OLM cells, we investigated metabotropic neuromodulation of intrinsic properties of OLM cells.

We found that both mAChR and mGluR activation increased the spontaneous action potential rate and caused the cells to exhibit long-lasting depolarizing plateau potentials following evoked spike trains.

Both the mAChR- and mGluR-induced increased spontaneous firing rate and plateau potentials were dependent on intracellular calcium, and were eliminated by blocking Ca^2+^-dependent transient receptor potential (TRP) cation channels. At the receptor level, Group I mGluRs were found to be responsible for the glutamatergic modulation of the plateau potentials. There was also a pronounced synergy between the cholinergic and glutamatergic modulation of the plateau potentials.

Our findings provide insights into how OLM cells are modulated by different neurotransmitters, and are likely to have functional implications on how OLM cells regulate hippocampal information processing during different brain states.

## Introduction

Hippocampal CA1 pyramidal cells receive two different streams of input: an input from the entorhinal cortex (EC) conveying a representation of the present external world, and an input from the hippocampal CA3 region conveying mnemonic information relating to past experiences ^1^.

Oriens-lacunosum moleculare (OLM) cells are inhibitory interneurons located in theCA1 region, and their activation has been implicated in regulating information flow in the hippocampus, facilitating CA3 input whilst diminishing EC input ^2^.

The CA1 network shifts rhythmically between responding to one or the other stream of input ^3^. Modulation of OLM cell activity may be a key contributor to this switching between functional modes in the hippocampus. However, the mechanisms underlying the modulation of OLM cells are poorly understood.

OLM cells somata are located in the oriens-alveus layer of the CA1, with dendrites that branch within the oriens layer where they receive glutamatergic input from local CA1 pyramidal cells ^4^ and cholinergic input from the medial septum ^2^. Their axons project to the lacunosum-moleculare layer, where they mediate pre- and post-synaptic inhibition of input to the distal apical dendritic tufts of hippocampal pyramidal cells and disinhibition to pyramidal cell proximal dendrites ^2,5,6^. OLM cells are characterised by spontaneous spiking in slice preparations ^2^ as well as in anaesthetised ^7^ and awake ^8^ animals, which implies that OLM cells provide their targets with a constitutive tonic inhibition ^9,10^.

OLM cells express both metabotropic cholinergic (mAChR) ^11-13^ and glutamatergic (mGluR) ^14,15^ receptors. Metabotropic effects have been previously reported in relation to LTP induction in OLM cells ^12,14^. However the effects of mGluR and mAChR modulation on the intrinsic properties of the cells and their responses to input (which may underlie these changes in plasticity) have not been fully addressed. Of particular interest is convergence of modulatory pathways in regulating neuronal behaviour, as shown in CA1 pyramidal cells ^16^. This is likely to occur often during arousal and wakefulness in behaving animals, when multiple neurotransmitter systems are active simultaneously ^17^, but which has so far not been assessed in other cell types.

Plateau potentials are extended periods of depolarisation following but in the absence of excitatory synaptic input, often caused by the opening of calcium channels ^18^. They are associated with LTP induction ^19^ and have been proposed as an underlying mechanism for working memory ^20^. Plateau potentials extend the period a post-synaptic cell can fire action potentials in response to excitatory input beyond the decay of the initial synaptic barrage. This creates an extended window where plasticity may be induced in a Hebbian ^19^ or non-Hebbian ^21^ manner, at behaviourally relevant time scales. mAChRs and mGluRs have been shown to cause the depolarising spike afterpotentials ^22,23^ and spontaneous and persistent firing ^24,25^ features associated with plateau potentials in a range of cell types across a variety of animal models.

Convergent evidence indicates that mGluRs are highly important for circuit functions in regards to conveying both semantic and modulatory information ^26^. Furthermore, both Group I and Group II mGluRs can be activated already by brief trains of 2-3 presynaptic spikes ^27^, indicating that only sparse activation is needed to elicit metabotropic effects. Since the glutamatergic synapses onto OLM cells are facilitatory ^28,29^, they probably belong to the so-called “Class 2” inputs ^27^, which are known to activate mGluRs in addition to ionotropic GluRs (iGluRs), suggesting that metabotropic activation is likely a regular feature at excitatory synapses on OLM cells.

Since OLM cells are probably crucially involved in several functions that are subject to neuromodulation, and these cells are themselves equipped with a variety of receptors and ion channel types that can mediate neuromodulation, it is highly relevant to explore this dimension of their functional repertoires. However, there have so far only been only few studies of neuromodulation in these cell types. This study aims at filling some of these gaps, by focusing on the neuromodulatory effects of two of the main neurotransmitters in the brain: glutamate and acetylcholine - each alone and in combination.

To study glutamatergic and muscarinic metabotropic modulation in hippocampal OLM cells we used a *Chrna2*-cre transgenic mouse line which has been shown to specifically expressed in OLM cells ^30^. We show that genetically identified OLM cells exhibit plateau potentials in response to both mAChR and mGluR activation. Pharmacological tests indicated that these responses were dependent on non-specific cation channels and the glutamatergic plateaus were mediated by group I mGluRs. mGluR activation also increased the spontaneous firing rate of the cells and increased excitability in response to current input. Our results suggest that metabotropic receptors activation of OLM cells dramatically increases their activity in response to excitatory input and allows ongoing action potential firing beyond the offset of depolarising inputs. We also suggest that intracellular signalling pathways of both mAChRs and mGluRs is convergent and simultaneous activation elicits synergistic supra-linear effects on plateau potential generation.

## Methods

### Ethical approval

All animal procedures were approved by the responsible veterinarian of the institute, in accordance with the statue regulating animal experimentation (Norwegian Ministry of Agriculture, 1996).

### Animals

*Chrna2-cre* transgenic C57BL6 mice were donated for breeding by the Kullander group and have been previously described ^2^. *Gt(ROSA)26Sor*^*tm14(CAG−tdTomato)Hze*^ (R26^*tom*^) transgenic mice were obtained from Jackson Laboratories. *Chrna2-cre* and *R26*^*tom*^ lines were crossbred to give a line expressing RFP variant tdTomato under control of the *Chrna2* gene (*Chrna2-cre; R26* ^*tom*^).

### Hippocampal slice preparation

Horizontal hippocampal slices were obtained from adult *Chrna2-cre; R26* mice. Mice were anaesthetized with isoflurane inhalation and decapitated, and the brain was removed quickly into ice-cold sucrose-based artificial cerebrospinal fluid (aCSF) containing (in mM): 1.25 NaCl, 1.25 KCl, 1.25 NaH_2_PO_4_, 7 MgCl_2_, 0.5 CaCl_2_, 16 glucose, 75 sucrose, 25 NaHCO_3_) saturated with 95% O_2_-5% CO_2_. 350 μm slices were obtained using a Leica VT1200 (Leica Microsystems) and incubated for 30 minutes at 35°C in aCSF containing (in mM): 125 NaCl, 2.5 KCl, 1.25 NaH_2_PO_4_, 1.4 MgCl_2_, 1.6 CaCl_2_, 16 glucose, 25 NaHCO_3_) saturated with 95% O_2_-5% CO_2_. After incubation slices were kept at room temperature until use.

### Electrophysiology

Whole-cell and cell-attached recordings were obtained using visual guidance from IR-DIC optics (BX51; Olympus, Tokyo, Japan) from the somata of OLM identified by widefield fluorescence microscopy.

Slices were maintained at32±0.5 °C and superfused with aCSF containing the AMPA, NMDA and GABA_A_ receptor blockers 6,7-dinitroquinoxaline-2,3-dione (DNQX; 10 μM), DL-2-Amino-5-phosphonopentanoic acid (_DL_-AP5;50 μM) and SR 95531 (gabazine; 5 μM) in order to block synaptic transmission. Pipettes (5-7 MΩ) were pulled from borosilicate glass tubing (outer diameter 1.5 mm, inner diameter 0.86 mm; Sutter Instruments, Novato, CA, USA) and filled with a solution containing (in mM): 120 potassium gluconate, 20 KCl, 5 phosphocreatine disodium salt, 4 MgATP, 0.4 NaGTP, 10 HEPES and 0.1 EGTA. The pH of the intracellular medium was adjusted to 7.2 with KOH, and osmolarity was between 280 and 290 mosmol^−1^. Recordings were made using a Multiclamp 700A (Molecular Devices; Sunnyvale, CA, USA), low pass filtered at 10 kHz, and digitized at 20 kHz. Access resistance was typically between 20 and 40 MΩ and was compensated at the beginning of every recording and adjusted as required.

### Data acquisition and analysis

Data was acquired using pCLAMP 10 and digitzed with a Digidata 1440 (Molecular Devices). Analysis was carried out using Clampfit (Molecular Devices), and results were plotted and statistical analysis performed in Origin 9.1 (OriginLab Corp; Northampton, MA, USA). Whilst recording current injection protocols, cells were held at a constant hyperpolarised baseline membrane potential of −60 mV by DC current injection. Cell-attached gap free recordings were made in voltage clamp with no external voltage command after obtaining a seal of 1 GΩ or tighter.

### Chemicals

DNQX, gabazine, DL-AP5, XE991, muscarine, t-ACPD, CPCCOet and MPEP were obtained from Tocris Bioscience (Bristol, UK). Potassium gluconate, flufenamic acid, and the other substances used for preparing the solutions were obtained from Sigma-Aldrich Norway AS (Oslo, Norway). All chemicals tested in the experiments were bath applied at a superfusion rate of ~2 ml min^−1^.

## Results

### Spontaneous firing in OLM cells is increased by muscarinic or metabotropic glutamate modulation

When recording from our genetically labelled cells (*n* = 14) in control conditions, we observed spontaneous firing of action potentials (3±0.74 Hz) in both cell attached voltage clamp (**Fig. 1Ai-iii**) and whole cell current clamp (**Fig. 1Bi-iii**) Such spontaneous firing has previously been reported in OLM cells in acute slice preparations ^2^, and we began by testing whether this was modulated by mAChR or mGluR activation. We observed a clear increase in the spontaneous action potential frequency during wash-in of either mAChR agonist muscarine or mGluR agonist tACPD (**Fig. 1C**). Thus, significant increases in firing were seen following application of either 10 μM muscarine (from 4.7 ± 1.0 Hz to 22.7 ± 3.3 HZ after muscarine application; *n* = 5; paired *t* test, *P*= 0.05; **Fig.1Di**) or 15 μM t-ACPD (from 0.6±0.4 to 23.6±7.1 Hz; *n* =6; *P* =0.02; **Fig. 1Dii**).

Both mAChR and mGluR activation are known to reduce the M-type potassium current (*I*_M_, mediated by Kv7 channels) activity ^31,32^, and modulation of these channels was found to affect resting membrane potential and spontaneous firing frequency in OLM cells ^15^. Therefore we sought to determine whether the increase in spontaneous firing seen in OLM cells during activation of these receptors was due to their blocking effect on Kv7 channels. Using the selective Kv7 channel blocker XE991 (10 μM), we observed significantly increased spontaneous firing compared to control conditions (from 4.7±2.6 to 11.1±5 Hz; *n* =14; *P* =0.04; **Fig. 1F**). Following application of XE991, either muscarine or t-ACPD was added to test whether their effects were occluded by prior Kv7 blockade. However, we observed no signs of such occlusion. Thus, addition of these agonists was followed by significant additional increases in spike frequencies, increases even greater than the difference observed between control and XE991 conditions. Addition of muscarine increased the spike frequency from 9.7±3.7 to 23.7±5.3 Hz (*n* =11; *P* =0.00005; **Fig. 1Fi**), and addition of t-ACPD increased the frequency from 11.1±5.1 to 38.2±11.2Hz (*n* =6; *P* =0.03; **Fig. 1Fii**). This suggests that the increased spontaneous firing seen with activation of mGluR and mAChRs is partly due to block of Kv7 channels, but that the increase is largely due to other mechanisms.

**Figure 1.**
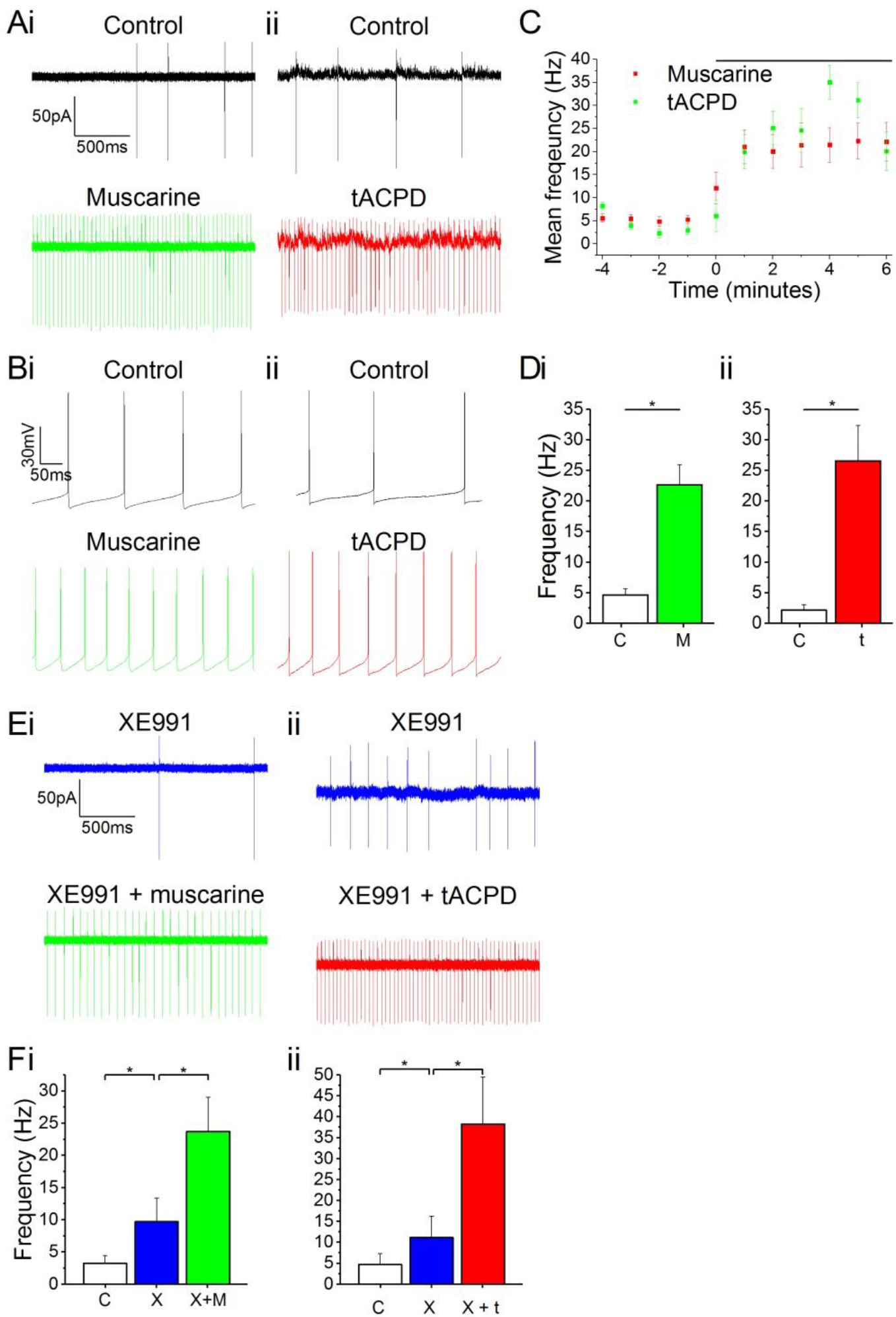
Increased spontaneous firing rate following mAChR/mGluR activation is only partly mediated by M current. ***A*** and ***B***, Example traces showing spontaneous action potential firing observed during gap free recording in control conditions in both cell attached (***A***) and whole cell **(*B***) patch configurations. The frequency of firing increased in response to application of either 10μM muscarine (***Ai* and *Bi***) or 15μM t-ACPD (***Aii*** and ***Bii***) ***C***, The rate of spontaneous firing during cell attached recording in control conditions and the increase in frequency following the application of either muscarine or tACPD (presence of drug indicated by black bar). Error bars = SEM. ∗p < 0.05 ***D***, Summary plot of the spontaneous firing frequency change for both muscarine (***Di***; *n* = 5) t-ACPD (***Dii***; *n* = 6) ***E*,** Example traces showing an increase in spontaneous firing following the application of either muscarine (***Ei***) or tACPD (***Eii***) in addition to XE991 ***F***, Summary plot of the spontaneous firing frequency change following XE991 application, and application of either muscarine (***Fi***; *n* = 7) or t-ACPD (***Fii***; *n* = 6). Kv7 blockade increased firing rate from control and application of mGluR or mAChR agonists significantly increased the firing rate further. Error bars = SEM. ∗*P* < 0.05.

### Plateau potentials in OLM cells

Lawrence et al. (2006) found plateau potentials in morphologically identified OLM cells following muscarinic activation, and we attempted to verify this finding in our genetically identified OLM cells. In line with previous reports, we observed that bath application of muscarine (10 μM) led to a change in post-pulse potential following intracellular injection of a 1 s long 50 pA depolarizing current pulse from a −60 mV holding potential: a shallow, slow afterhyperpolarization (AHP) in the normal extracellular medium (control) was replaced by a slow depolarising plateau afterpotential accompanied by continued spiking, typically lasting more than a second (**Fig. 2Ai**). Application of tACPD (15 μM) produced a similar effect (**Fig. 2Aii**). Similar to previous studies, we defined the post-burst potential as the difference between the mean membrane potential during a 200 ms window prior to the current pulse (dotted lines in **Fig. 2 A-B**) and the mean membrane potential during a 200 ms time window following 100 ms after the offset of the current pulse injection ^15^. The post-pulse potential changed significantly from control following application of either muscarine (from −2.6±0.4 mV to 8.2±3.4 mV; *n* = 6; *P* = 0.02; **Fig. 2Ci**) or tACPD (from −2.8±0.7 mV to 7.7±2.5 mV; *n* = 6; *P* = 0.008; **Fig. 2Cii**).

**Figure 2.**
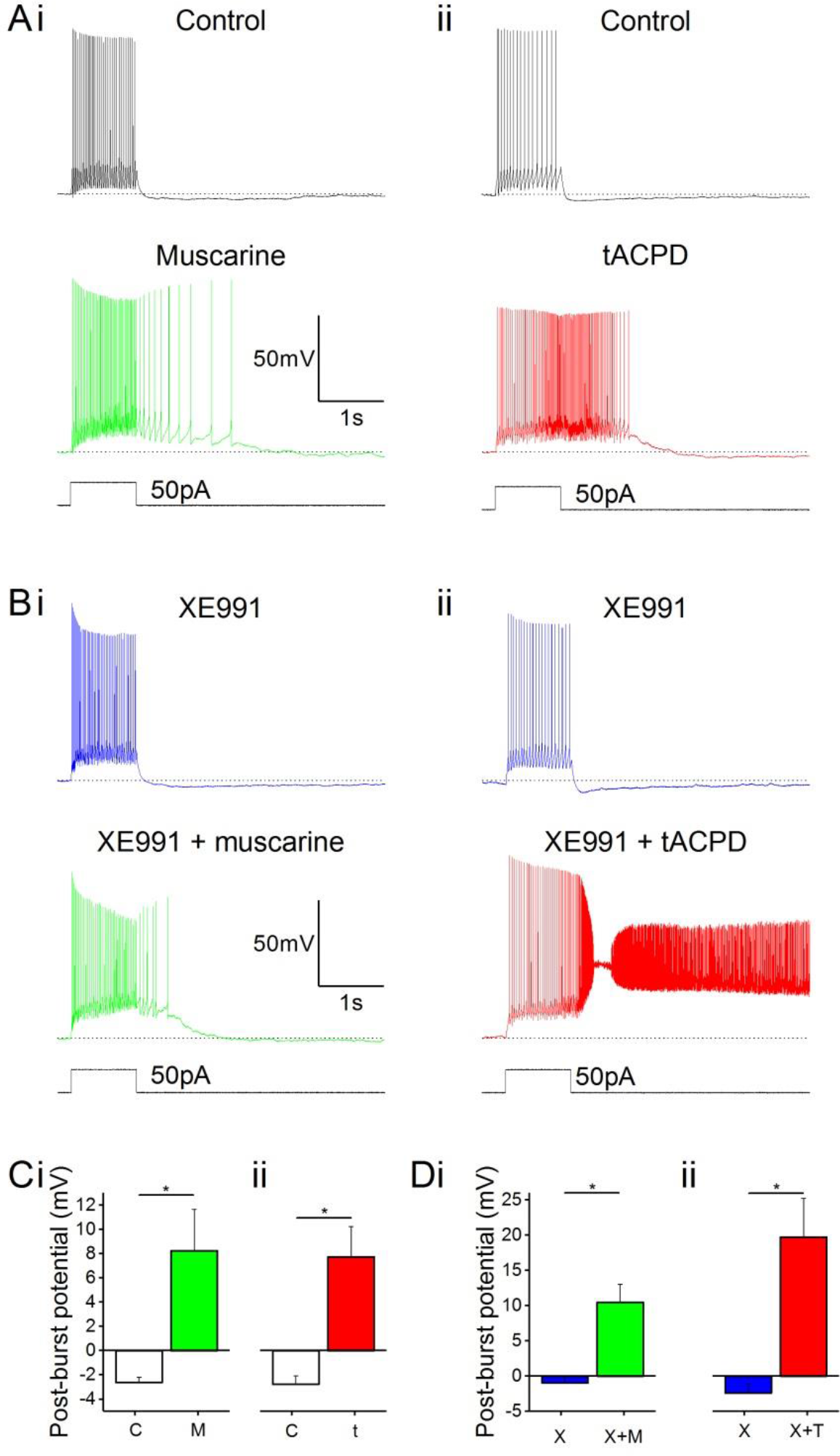
mAChR and mGluR activation induce plateau potentials in OLM cells. ***A***, Example traces showing the onset of a plateau potential following the application of either muscarine (***Ai***) or t-ACPD (***Aii***) ***B***, Example traces showing that blocking M-current with XE991 did not induce post-pulse spiking or plateau potentials but could be induced with additional application of muscarine (***Bi***) or tACPD (***Bii***). The horizontal, dotted lines in ***A-B*** indicate the baseline membrane potential during a 200 ms time window prior to the current pulse onset ***C***, Summary plots showing the change in post-burst potential from control with either application of either muscarine (***Ci***; *n* = 6) or tACPD (***Cii***; *n* = 7), moving from an AHP to an afterdepolarisation (ADP) in both conditions ***D***, Summary plots showing the change in post-burst potential in the presence of XE 991 and following the application of either muscarine (***Di***; *n* = 6) or t-ACPD (***Dii*;** *n* = 6). Both agonists induced a significant change in the post-pulse potential following in the presence of XE 991. Error bars = SEM. ∗*P* < 0.05.

### Modulation of the M current is not responsible for the changes in post-burst potential following muscarinic or glutamatergic metabotropic modulation

Having observed that both muscarinic or glutamatergic metabotropic receptor activation induce a depolarizing plateau potential in the OLM interneurons, we searched for the underlying ionic mechanism.

*I*_M_, which is reduced by both mAChR and mGluR activation ^31-33^, is known to contribute to afterhyperpolarizations (AHPs) in mammalian central neurons both directly ^34^ and indirectly. Thus, a metabotropic suppression of the hyperpolarizing *I*_M_, might possibly cause or unmask a depolarizing plateau. To test this hypothesis, we bath-applied the selective *I*_M_ blocker XE991 ^35^ and compared the post-burst potential before and after adding either muscarine (**Fig. 2Ai, Bi**) or tACPD (**Fig. 2Aii, Bii**), in the absence (**Fig. 2A**) or presence (**Fig. 2B)** of 100 μM XE991, in order to determine the contribution of M-current to the post-burst potential. Shallow AHPs were still observed in the presence of XE991 (**Fig. 2Bi, Bii**), and significant changes with emergence of large plateau potentials were still seen (like without XE991) with addition of either muscarine (from −1.0±0.1 mV (AHP) to 10.5±2.6 mV (plateau); *n* = 6; *P* = 0.006; **Fig. 2Di**) or tACPD (from −2.4±1.3 (AHP) to 19.8±5.6 mV (plateau); *n* = 6; *P* = 0.008; **Fig. 2Dii**). This showed that plateau potentials can still be generated in these cells during blockade of *I*_M_. Therefore another mechanism besides *I*_M_ modulation is necessary to explain the induction of plateau potentials by mGluR and mAChR activation.

### Calcium dependence of glutamatergic and muscarinic plateau potentials

Having excluded that the depolarizing plateau potentials are merely due to suppression of the hyperpolarizing *I*_M_, we tested other possible mechanisms. Since several neuronal afterpotentials are calcium dependent, including muscarine-induced plateaus in stratum oriens interneurons ^15^, we tested whether reducing either extracellular or intracellular [Ca^2+^] affected the plateau potentials in OLM cells in response to muscarine and tACPD.

After switching from our standard control aCSF extracellular medium to a Ca^2+^-free medium in the constant presence of either muscarine (**Fig. 3Ai)** or tACPD **(Fig. 3Aii)**, we observed a significant reduction in the post-burst plateau potentials in the presence of either muscarine (from 9.6±8.6 (plateau) to −1.±3.4 mV (AHP); *n* =7; *P* =0.02; **Fig. 3Ci**) or tACPD (from 8.0±3.6 to (plateau) - 1.1±2.3 mV(AHP); *n* =6; *P* =0.04; **Fig. 3Cii**), the trend moving from a plateau to an AHP in both cases.

We next tested the effects of reducing the intracellular calcium level by including 10 mM of the calcium chelator BAPTA in the pipette solution. The plateau potentials were observed immediately after break-in (**Fig. 3Bi**,0 min.), but gradually disappeared as BAPTA diffused into the cell from the pipette, still in the continuous presence of muscarine in the bath (**Fig. 3Bi**) or tACPD (**Fig. 3Bii**). Thus, after 10 minutes the post burst plateau potential was significantly reduced, both in the presence of muscarine (from 10.1±4.6 to 3.2±2.4 mV; *n* =5; *P* =0.04; **Fig. 3Di**) and with tACPD (from 27.7±2.9 to 0.4±0.8 mV; *n*=8; *P* =0.00001; **Fig. 3Dii**). The presence of clear afterhyperpolarizations (AHPs) in Ca^2+^-free medium and with BAPTA in at least some cells (Fig. 3Ai and ii, lower panels), indicates that these rather slow AHPs are not caused by Ca^2+^-dependent potassium currents, unlike slow AHPs in many vertebrate principal neurons ^36^.

**Figure 3.**
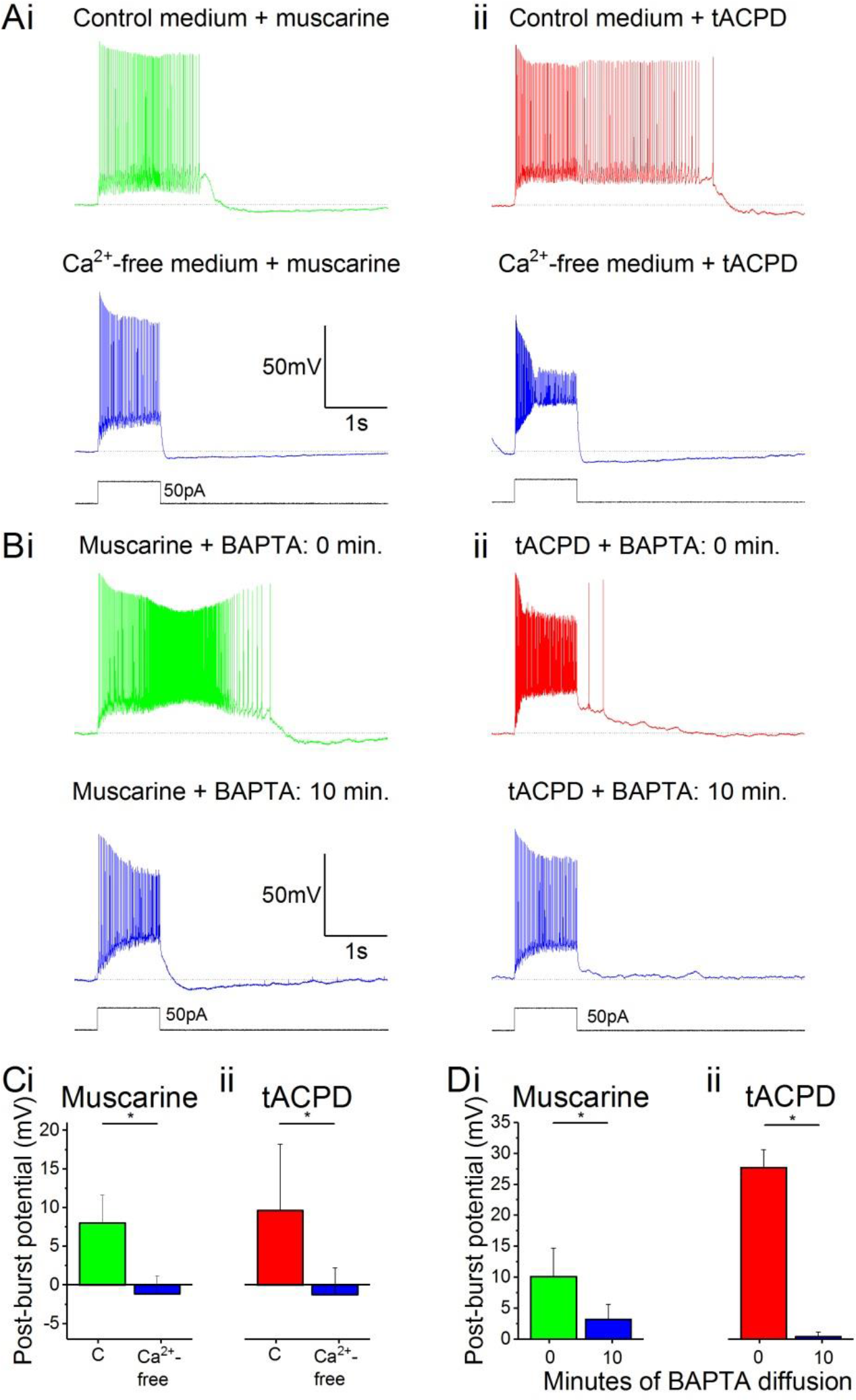
The muscarinic and glutamatergic plateau potentials are calcium-dependent. ***A***, Example traces of the abolition of muscarine- (***Ai***) or tACPD-induced (***Aii***) plateau potentials by switching to a Ca^2+^-free aCSF medium in the continuous presence of agonists ***B***, Example traces showing the abolition of muscarine- (***Bi***) or tACPD-induced (***Bii***) plateau potentials immediately after (upper traces) and 10 minutes after break in (lower traces) when the BAPTA-containing pipette solution became contiguous with the intracellular space ***C***, Summary plots of the change in post-burst potential in the presence of either muscarine (***Ci***; *n* = 6) or tACPD (***Cii***; *n* = 7) after switching to a Ca^2+^-free medium, showing a move from an ADP to an AHP ***D***, Summary plots of the change in post-burst potential in the presence of muscarine (***Di***; *n* = 5) or tACPD (***Dii***; *n* = 8) after 10 minutes of allowing BAPTA to diffuse into the intracellular solution. Error bars = SEM. ∗*P* < 0.05.

### TRPC channels mediate plateau potentials induced by mAChRs and mGluRs

We next asked what channel could underlie the calcium entry we previously showed to be essential for the generation of plateau potentials induced by mAChR or mGluR. A plausible candidate mechanism is transient receptor potential C (TRPC) channel conductances which have been shown to mediate plateau potentials in cortical pyramidal cells ^37^. To test this idea, we used non-specific TRPC channel blocker flufenamic acid (FFA) following application of either mAChR or mGluR agonists.

Application of 50 μM FFA blocked the plateau potentials induced by 10 μM muscarine (**Fig. 4Bi**) and 15 μM tACPD (**Fig. 4Aii**). In both cases, the post-pulse potentials were significantly reduced by FFA, from 4.2±1.4 to 0.9±0.5 mV after muscarine (*n* = 5; *P* = 0.002; **Fig. 4Bi**), and from 24.9±4.3 to 1.9±1.0 mV or tACPD (*n* = 4; *P* = 0.007; **Fig. 4Bii**).

**Figure 4.**
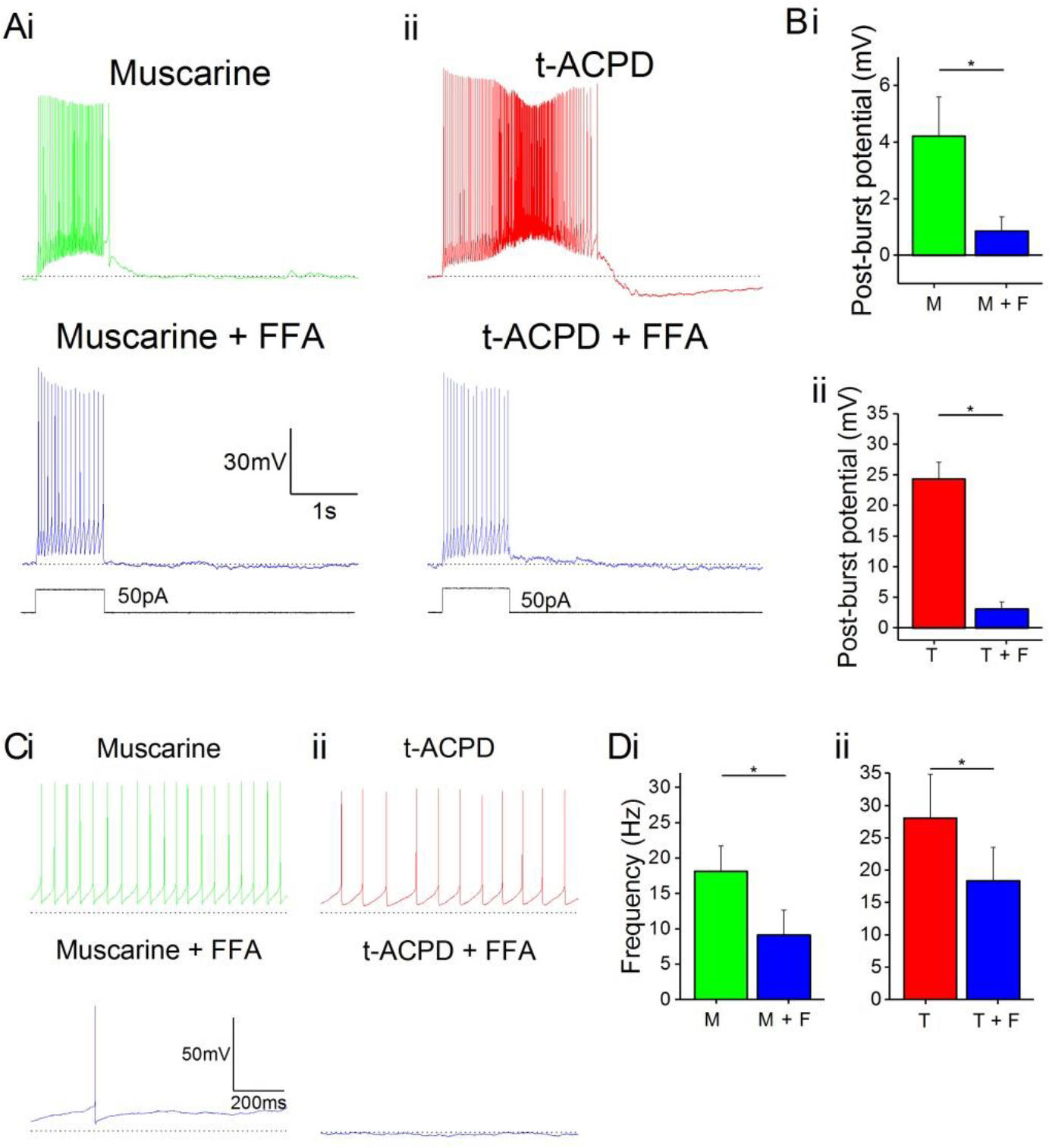
mAChR and mGluR agonist-induced effects can be reversed by non-specific TRP inhibitor. ***A***, Example traces showing plateau potentials induced (upper voltage traces) by application of 10 μM muscarine (***Ai***) or 15 μM tACPD (***Aii***) are blocked by addition of FFA (lower voltage traces) ***B***, Summary plots quantifying post-pulse potential following application of FFA in the presence of either muscarine (***Bi***; *n* = 5) or tACPD (***Bii***; *n* = 4). In both cases it was significantly reduced ***C***, Example traces showing spontaneous firing in the presence of muscarine (***Ci***) or tACPD (***Cii***) before (upper trace) and after (lower trace) the addition of FFA ***D***, Summary plots of the spontaneous firing rate following application of FFA in the presence of either muscarine (***Di***; *n* = 10) or tACPD (***Dii***; *n* = 10). Error bars = SEM. ∗*P* < 0.05. It significantly decreased in the presence of either mGluR or mAChR agonists.

FFA also reduced the spontaneous firing in the presence of either muscarine (**Fig. 4Ci**) or tACPD (**Fig. 4Cii**). Spontaneous action potential frequency was significantly reduced by FFA both when applied in the presence of either muscarine (from 18.2±3.6 to 9.2±3.5 Hz; *n* = 10; *P* = 0.002; **Fig. 4Di**) or tACPD (from 28.1±6.7 to 18.4±5.2 Hz; *n* = 6; *P* = 0.04; **Fig. 4Dii**).

### Group I mGluRs are responsible for the glutamatergic plateau potentials

tACPD is an agonist at both group I and group II mGluRs ^38^. Hence, we sought to determine if activation of one group was more responsible than the other for the phenomena of plateau potential induction and increased spontaneous firing. By applying antagonists for group I receptor subtypes mGlu1 Rs (CCPOET; 100 μM) and mGlu5 Rs (MPEP; 60 μM) in the presence of tACPD, we observed that plateau potentials were abolished (**Fig. 5A**) and the post-burst potential significantly decreased (from 22±5.5 to 3.4±2.9 mV; *n* = 6; *P* = 0.02; **Fig. 5Ei**).

**Figure 5.**
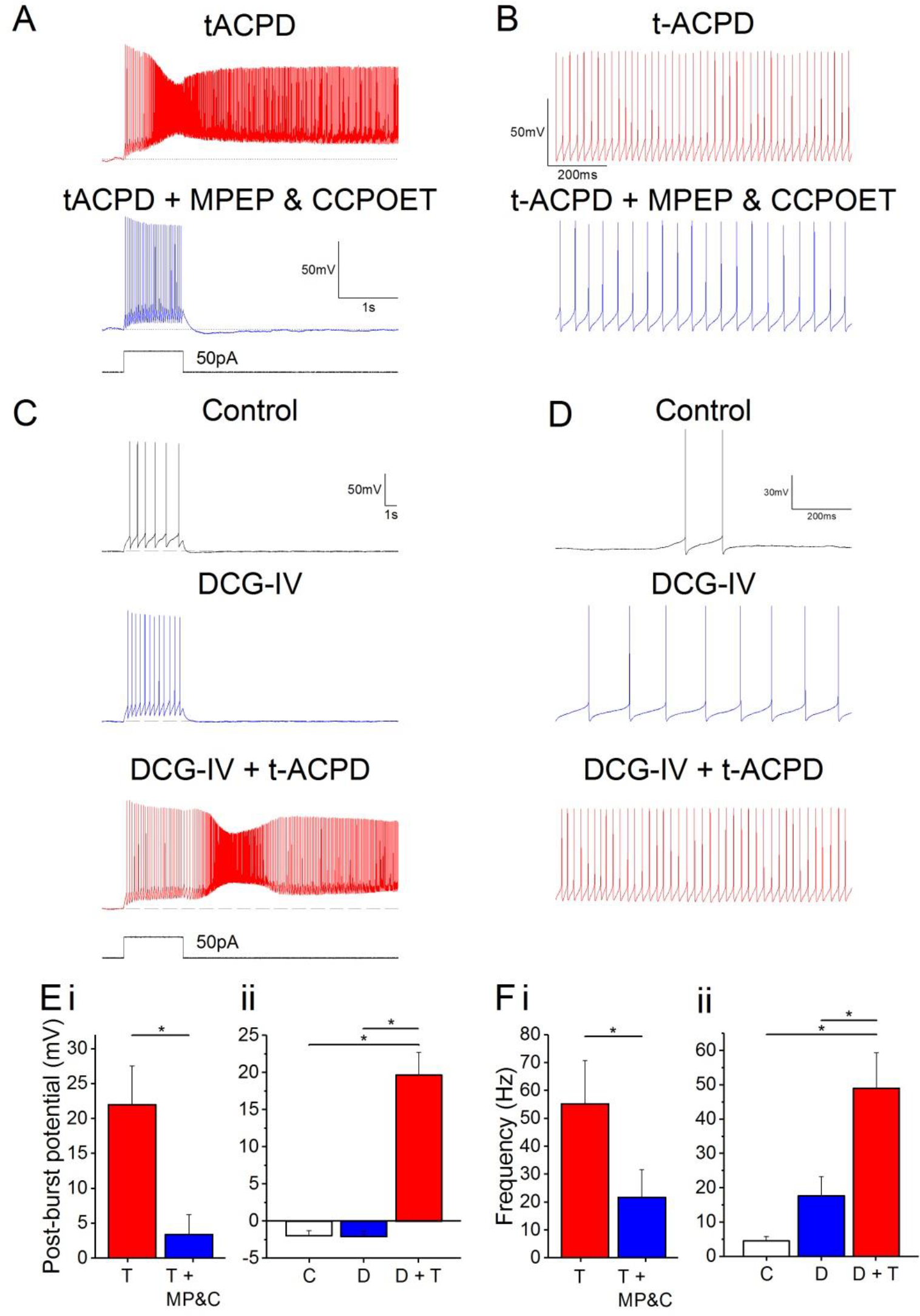
Plateau potentials evoked by group I but not group II mGluR activation. ***A***, Example traces of response to 50 pA 1s current pulse in the presence of 15 μM tACPD, before and after application of MPEP and CCPOET ***B***, example traces showing the decrease in spontaneous firing in the presence of t-ACPD following addition of MPEP and CCPOET ***C***, Example traces of plateau potentials following application of DCG-IV and DCG-IV & tACPD ***D***, Example traces of spontaneous firing following application of DCG-IV and DCG-IV & tACPD ***E***, Summary plot of the post-pulse potential in the presence of t-ACPD following the addition of MPEP and CCPOET (***i***; *n* = 5) and following DCG-IV application and subsequence tACPD application (***ii***; *n* = 5) ***F***, Summary plot of spontaneous firing frequency in the presence of tACPD following the application of MPEP and CCPOET (***i***; *n* = 4), and in response to DCG-IV and following sequential application of tACPD (***ii***; *n* = 5). Error bars = SEM. ∗*P* < 0.05.

These mGlu1 and mGlu5 R antagonists also produced a change in the spontaneous firing in the presence of tACPD (**Fig. 5B**). Firing frequency was significantly reduced (from 55.2±15.6 to 21.7±10 Hz; *n* = 4; *P* = 0.02; **Fig. 5Fi**), but did not reach the previously established levels of spontaneous firing in control conditions (0.57651±0.36898 Hz; **Fig. 1D**).

These results indicated that the tACPD-induced plateau is wholly dependent on group I mGluR activation whilst the tACPD induced increased in spontaneous firing is at least partially dependent of group I mGluRs. To test the impact of group II mGluRs, we added a selective agonist (DCG-IV ^39^; 15 μM) in the absence of tACPD. The post-burst potential did not change with application of DCG-IV (from −2.0±0.7 to −2.1±0.8 mV; *n* = 5; *P* > 0.05; **Fig. 6C**) but subsequent application of tACPD in the same cells elicited a significant increase (from −2.1±0.8 to 19.6±3.1 mV; *n* = 5; *P* = 0.03; **Fig. 5C**) as expected from the previous experiments (**Figures 1-4**).

Spontaneous firing appeared to change with application of DCG-IV (**Fig. 5D**), but was not statistically significant (from 4.5±1.3 to 17.7±5.6 Hz; *n* = 7; *P* > 0.05; **Fig. 5Fii**) whilst the increases in frequency caused by application of tACPD were shown to be significant (from 17.7±5.6 to 49±10.4 Hz; *n* = 7; *P* = 0.03; **Fig. 5Fii**).

**Figure 6.**
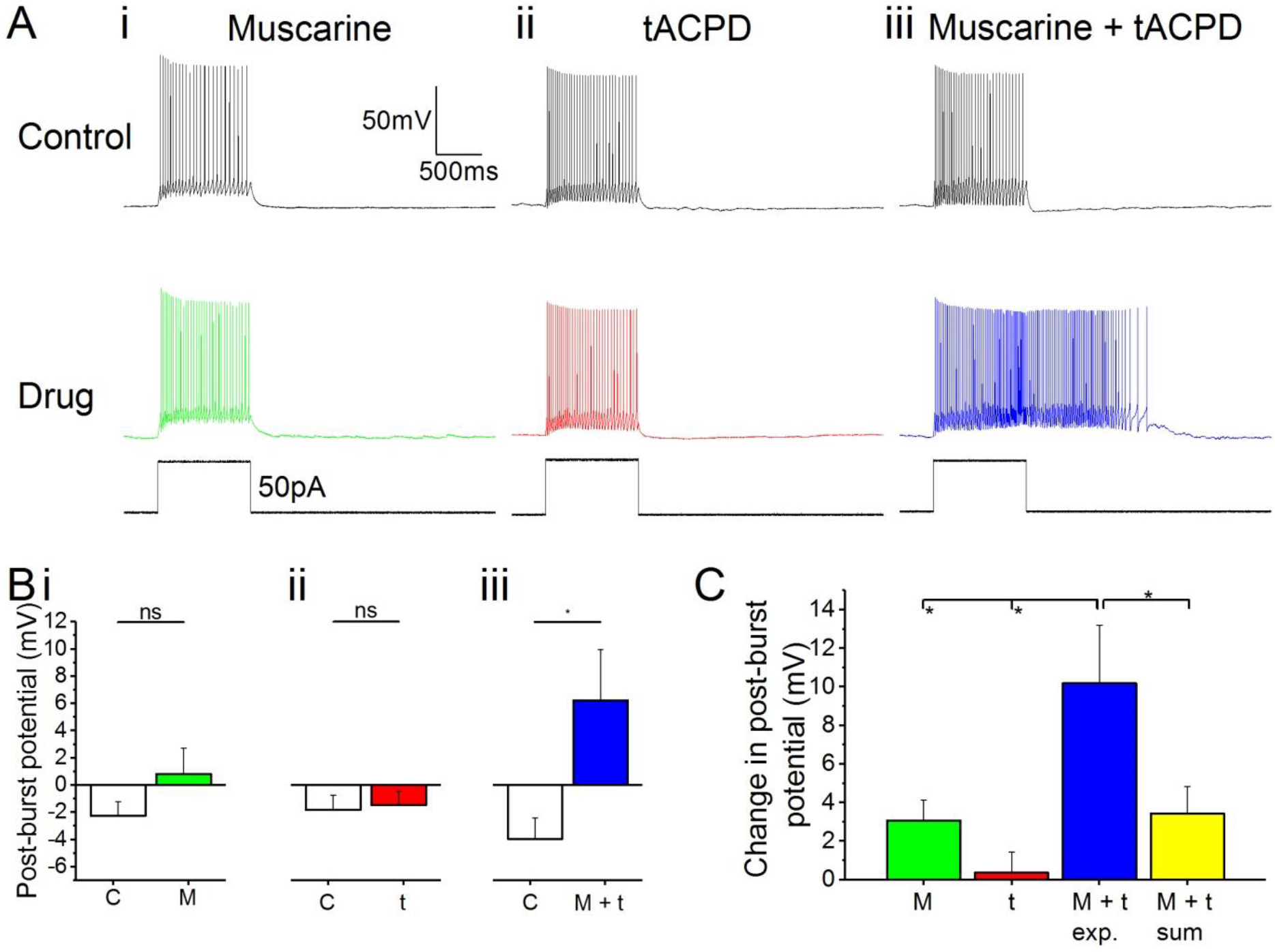
Synergistic effects on post-pulse potential by coapplication of low doses of muscarine and tACPD. ***A***, Example traces of OLM cells in response to a 50 pA 1 s current pulse before and after application of either low dose muscarine (***Ai***; 4 μM; *n* = 8), low dose tACPD (***Aii***; 4μM; *n* = 8), or both (***Aiii***; 4 μM + 4 μM; *n* = 6) ***B***, Summary plots show no significant change in post-burst potential was seen with application of low dose of muscarine (***Bi***; *n* = 8) or tACPD (***Bii***; *n* = 8), however a significant increase was seen following application of a low dose of both drugs simultaneously (***Biii***; *n* = 6), moving the post-burst potential from and AHP to ADP ***C***, Summary plot for change in post-pulse potential, in all experimental conditions and also post-hoc linear sum (yellow column). The linear sum was calculated from 8 randomly selected pairs of independent measures of single low dose drug administration. Error bars = SEM. ∗*P* < 0.05. ns, not significantly different, *P* > 0.05.

### Non-linear synergistic induction of plateau potentials by mAChR and mGluR co-activation

In many *in vivo* situations, such as in awake, behaving animals, it is likely that both cholinergic and glutamatergic metabotropic neuromodulation may occur simultaneously in the same cell, activated by distinct inputs via their respective receptors. It is then functionally important whether these converging forms of metabotropic neuromodulation are occluding each other, or are linearly additive, or perhaps non-linearly synergistic. To explore this issue, we used low concentrations of the agonists (4 μM muscarine; 4 μM tACPD), applied both individually and simultaneously. We observed no plateau potentials after adding either the low dose of muscarine alone (**Fig. 6Ai**) or after adding the low dose of tACPD alone (**Fig. 6Aii**), but simultaneous application of both these agonists led to an emergence of a plateau (**Fig. 6Aiii**). This was confirmed by the lack of significant change in post-burst potential in the presence of either low dose muscarine (from −2.3±1 to 0.8±1.9 mV; *n* = 8; *P* > 0.05; **Fig. 6Bi**) or low dose tACPD (from −1.8±1.1 to −1.5±1 mV; *n* = 8; *P* > 0.05; **Fig. 6Bii**), but a significant increase in the presence of both (from −3.9±1.5 to 6.2±3.7 mV; *n* = 6; *P* = 0.01; **Fig. 6Biii**), which caused the transformation of the post-burst potential from an AHP to an ADP.

To determine whether the resultant effect of combining low doses experimentally was any different from what might be expected from the addition of effects seen in the presence of individual drugs; we summed 8 randomly selected pairs of measurements from independent drug administrations and compared the sum with the mean values obtained during experimental recordings in the presence of both agonists.

The mean change in post-burst potential in the presence of both muscarine and tACPD (**Fig. 6C;** 10.2±3 mV; *n* = 8) was significantly increased from control when compared to the values obtained in the presence of either just muscarine (3.1±1.1 mV; *n* = 8; *P* = 0.02) or just tACPD (0.4±1.13 mV; *n* = 8; *P* = 0.004). When the low concentrations of agonists were applied together, the post-burst potential was significantly greater than the sum of the effects of low concentrations of agonists applied separately (10.2±3 simultaneous versus 3.4±1.4 mV summed; summed *n* = 8; *P* = 0.04).

## Discussion

Our results show that OLM cell activity and response to synaptic input is highly modulated by glutamatergic and cholinergic neuromodulatory systems. We identify the receptor types responsible for the modulation and the downstream signalling targets of these receptors which lead to the modulatory changes. Finally we observe pronounced synergy between the cholinergic and glutamatergic branches of the modulation, suggesting an integral role for these cells in switching information flow in the CA1 circuit; between functional modes of the hippocampus; and potentially between different brain states.

### Modulation of spontaneous firing

We found that OLM cells increased their rate of spontaneous firing in response to agonists of either metabotropic acetylcholine receptors or metabotropic glutamate receptors. Depolarization and increased frequency of spontaneous firing have been shown in multiple cell types in response to mAChR ^40,41^ or mGluR ^23,42^ activation. In OLM cells, cholinergic and glutamatergic metabotropically induced increase in firing rate will enhance the tonic inhibitory control over their postsynaptic targets: increasing inhibition to the distal apical dendritic compartment of pyramidal cells ^6^ whilst disinhibiting the proximal apical dendrite ^2^. This will prioritise the proximal Schaffer collateral CA3 inputs over the distal performant path EC inputs. OLM cell activity has been suggested as a mechanism for supressing current, incoming sensory information from the EC and thereby instead promoting the processing of afferent information from the CA3, where learnt patterns are stored in the autoassociative CA3 network ^2^. This may suggest that conditions that increase OLM cell activity may tend to impair memory acquisition and promote memory retrieval ^43^.

Increased OLM cell activity in response to mAChR activation could underlie the observation that activation of the medial septum, which provides cholinergic inputs to OLM cells during wakefulness and arousal, impaired new memory formation in rats ^44^ whilst inactivation diminished memory persistence ^45^.

OLM cells receive facilitating excitatory input, suggesting they have “class 2” glutamatergic synapses which contain both iGluRs and mGluRs ^27^. They also express nicotinic AChRs ^46^, and of course iGluRs. This suggests that the depolarization mediated by glutamatergic or cholinergic input will occur via both ionotropic and metabotropic receptors, and hence the level of depolarisation will be even greater than we observed here.

The longer duration of metabotropic compared to ionotropic receptor-mediated postsynaptic effects suggests that increases in spontaneous firing may persist long after the decay of the fast synaptic events, leading to longer periods of higher frequency spontaneous firing.

Whilst we saw some increase in spontaneous firing in response to blocking the M-current as would be expected ^47^, subsequent addition of either mAChR or mGluR agonists elicited a much larger, further firing frequency increase. The magnitude of this increase suggests that the mAChR or mGluR activation-dependent increase in spontaneous firing rate is mediated primarily through another mechanism, though parallel modulation of *I*_M_ may also play a role.

Previous work has shown that TRP channels are activated by mGluRs ^48,49^ and mAChRs ^50^. We saw that the increase in spontaneous firing caused by mAChR or mGluR activation was reduced in the presence of the TRP-channel blocker FFA, suggesting that TRP channels play an important role in regulating the prevailing membrane potential and the spontaneous firing frequency in these cells.

### Depolarizing plateau potentials

mAChR agonists have been previously shown to induce plateau potentials in morphologically identified putative-OLM cells ^15^ and we confirmed that this occurs in OLM cells.

Acetylcholine-dependent plateaus in OLM cells were shown to be mediated by type 1 and type 3 mAChRs ^15^, both of which associate with G-proteins of the G_q_ type ^51^. We found that the mGluR subtypes responsible for glutamate-dependent plateaus were group I mGluRs; this group includes mGluR_1_ and mGluR_5_ both of which also associate with G_q_ ^52^. This indicates that G_q_ is probably responsible for inducing plateau potentials in OLM cells. Interestingly, in spinal cells, G_q_-coupled 5-HT2 receptors also promote plateau potentials ^53^, further suggesting the crucial involvement of G_q_ across neuron types, and suggesting that other G_q_-coupled receptors present in OLM cells may also induce or contribute to plateau potentials.

Plateau potentials are associated with a depolarization of the membrane potential, accompanied not just by increased spiking but also by burst of spikes ^54^. These bursts may signal to downstream targets of OLM cells that a specific convergence of signals of distinct origins or transmitter systems has occurred.

Plateau potentials in cortical cell dendrites are known to facilitate LTP induction during sensory processing ^19^, as the prolonged depolarization and spiking associated with plateaus increases the window of opportunity for Hebbian pairing. The plateau itself, even without additional spiking, may be enough to enhance synaptic transmission, and may underlie the observations of Bittner, et al. ^21^ of potentiation of input that arrived seconds before and after complex spiking (“behavioural time scale plasticity”) during learning. If such plasticity can also occur in OLM and other inhibitory interneurons, in addition to the excitatory cells where it has previously been observed, this may substantially expand the possibilities for plasticity and tuning of hippocampal and other cortical microcircuits during learning.

### TRP channels and calcium-dependence

We observed that plateau potentials were abolished by the specific TRP channel blocker flufenamic acid (FFA), indicating that they were dependent on inward calcium current through these channels. This conclusion is supported by our finding that the plateau potentials were also abolished by Ca^2+^- free medium and by intracellular BAPTA, indicating Ca^2+^-dependence. Interestingly, however, the plateau potentials we observed were NMDA-R-independent, whereas some plateaus in other cell types has been shown to be dependent on NMDA-R-mediated Ca^2+^-entry ^19^. Since CA1 OLM cells do not express NMDA receptors ^55^, but the plateau potentials we observed were nevertheless dependent on Ca^2+^ entry, it appears that the activation of TRP channels by mGluRs has functionally replaced the functional role played by Ca^2+^-permeable iGluRs (NMDA-Rs) in other cell types.

TRP-dependent plateau potentials have been observed in other cells types ^37^. In addition, voltage-gated calcium (Ca_V_) channels, activated downstream of metabotropic receptors, have been shown to generate Ca^2+^-dependent plateau potentials in different cell types, involving R-type ^16^, N-type ^56^, and L-type ^57,58^ Ca_V_ channels. This suggests that plateau potentials represent a conserved response feature implemented by multiple cell types employing different Ca^2+^-permeable ion channels present in their membranes.

We showed that reducing the free intracellular calcium level by buffering prevented the emergence of plateau potentials. This likely prevents activation of the Ca^2+^-dependent TRP cation channels, rather than preventing the triggering of release of Ca^2+^-induced Ca^2+^ release, which has been shown to be not involved in plateau potential generation elsewhere ^59^.

From the present study, we cannot determine precisely which member(s) of the TRP channel family are responsible for the observed plateau potentials, though based on known sensitivity to FFA of a range of TRP channels, some candidates can be suggested based on the concentration of FFA we used and saw to be effective (TRPC6, TRPM5,TRPM3, TRPC5, TRPV4 & TRPC4 have an IC50 at around the concentration of FFA we used, according to Guinamard, et al. ^60^). Since specific pharmacological blockers for only a few TRPC family members are available, future identification of the exact subtype would most likely require a transgenic knockout approach.

### mGluR subtypes

Previous work in pyramidal neurons has shown that group I mGluR activation leads to depolarization ^42,61^, whereas group II activation leads to hyperpolarization ^62^. Therefore, the depolarization we observed to be caused by the pan-mGluR agonist t-ACPD suggests that group I mGluRs are most likely to be the predominant mGluR subtype group underlying this type of modulation in OLM cells. We observed no significant increase in spontaneous firing in response to a group II mGluR agonist, and antagonists at group I mGluRs decreased the spontaneous firing frequency that had previously been increased by a pan-mGluR agonist, further suggesting that glutamatergic modulation of spontaneous firing of OLM cells is dominated by group I subtype receptors.

Whilst the increase in spontaneous firing did not reach statistical significance in our experiments (*P* = 0.08765), a clear trend of increase was observed which goes against previous observations of hyperpolarization in response to group II agonists in other cells ^62^.

### Synergy

Plateau potential generation also results from convergence of spatially segregated inputs. For example, Takahashi and Magee ^54^ showed CA1 pyramidal cells receiving simultaneous glutamatergic inputs from CA3 and EC exhibited NMDA-R-dependent plateau potentials in their distal dendrites. Their study also showed that the summation of inputs was multiplicative, suggesting that non-linear summation may be a characteristic feature of plateau potential generation, whether they are dependent on ionotropic receptors, or metabotropic receptors as our data shows.

Park and Spruston ^16^ showed synergistic effects on post-burst ADPs following mGluR and mAChR activation in CA1 pyramidal cells. Whilst these lasting depolarisations were also non-linear, they were not accompanied by additional spiking. This is possibly because they observed the ADP following only 5 spikes, whilst we generated many more with a square current step, and the size of the ADP/plateau potential is probably related to the number of preceding spikes ^63,64^.

Whilst it is striking that the receptors that are responsible for plateau potentials all seem to couple to the same G-protein, it is possible that there are also other intracellular effectors, since some G-protein coupled receptors also seem to act via pathways independent of their G-protein ^65^. In particular group I mGluR-mediated effects such as depolarization have been reported to be insensitive to inhibition of G-protein signalling ^66^. Therefore, future work is needed to identify the pathways responsible for plateau potentials and determine the mechanism underlying the supra-linear synergy that we observed here.

### Conclusion

Overall, this study has demonstrated the presence of plateau potentials in Chrna2 positive OLM cells in the CA1 field of the rodent hippocampus, in response to metabotropic glutamate or acetylcholine modulation, and shown the consequences of such modulation on intrinsic properties of these cells. Our results suggest that OLM cells respond as a coincidence detector of glutamatergic (probably cortical/hippocampal, and/or thalamic) and cholinergic (probably largely septal) input and produce supra-linear output to its targets where they play a vital role in controlling input priority to the hippocampus and in controlling memory encoding and recall functions.

## Acknowledgements

AcknowledgementsThis study was partially supported by the European Union’s Horizon 2020 research and innovation programme under grant agreement 7202070 (Human Brain Project (HBP)) and the Norwegian Research Council (NRC: 262950/F20 and 214079/F20).

